# Fiscore Package: Effective Protein Structural Data Visualisation and Exploration

**DOI:** 10.1101/2021.08.25.457640

**Authors:** Auste Kanapeckaite

**Affiliations:** Algorithm379; University of Reading, School of Pharmacy, Hopkins Building, Reading RG6 6UB, United Kingdom

## Abstract

The lack of bioinformatics tools to quickly assess protein conformational and topological features motivated to create an integrative and user-friendly R package. Moreover, the *Fiscore* package implements a pipeline for Gaussian mixture modelling making such machine learning methods readily accessible to non-experts. This is especially important since probabilistic machine learning techniques can help with a better interpretation of complex biological phenomena when it is necessary to elucidate various structural features that might play a role in protein function. Thus, *Fiscore* builds on the mathematical formulation of protein physicochemical properties that can aid in drug discovery, target evaluation, or relational database building. In addition, the package provides interactive environments to explore various features of interest. Finally, one of the goals of this package was to engage structural bioinformaticians and develop more robust R tools that could help researchers not necessarily specialising in this field. Package *Fiscore* (v.0.1.3) is distributed via CRAN and Github.

## Introduction

*Fiscore* R package was developed to quickly take advantage of protein topology/conformational feature assessment and perform various analyses allowing a seamless integration into relational databases as well as machine learning pipelines (Kanapeckaitė et al., 2021). The package builds on protein structure and topology studies which led to the derivation of the Fi-score equation capturing protein dihedral angle and B-factor influence on amino acid residues (Eq.1&2) (Kanapeckaitė; et al., 2021). The introduced tools can be very beneficial in rational therapeutics development where successful engineering of biologics, such as antibodies, relies on the characterisation of potential binding or contact sites on target proteins (Kanapeckaitė et al., 2021; Du et al., 2020). Moreover, translating structural data into scores can help with target classification, target-ligand information storage, screening studies, or integration into machine learning pipelines (Kanapeckaitė; et al., 2021; Du et al., 2020). As a result, Fi-score, a first-of-its-kind *in silico* protein fingerprinting approach, created a premise for the development of a specialised R package to assist with protein studies and new therapeutics development (Kanapeckaitė et al., 2021).

*Fiscore* package allows capturing dihedral angle and B-factor effects on protein topology and conformation. Since these physicochemical characteristics could help with the identification or characterisation of a binding pocket or any other therapeutically relevant site, it is important to extract and combine data from structural files to allow such information integration (Kanapeckaitė et al., 2021; Fauman et al., 2011; Faraggi et al., 2009). Protein dihedral angles were selected as they contain information on the local and global protein structural features where protein backbone conformation can be highly accurately recreated based on the associated dihedral angles (Kanapeckaitė et al., 2021; Faraggi et al., 2009). Furthermore, since Ramachandran plot, which provides a visualisation for dihedral angle distributions, namely *ϕ* (phi) and *ψ* (psi), allows only a holistic description of conformation and cannot be integrated with traditional parametric or non-parametric density estimation methods, a specific transformation was required to use this data. An additional parameter, specifically the oscillation amplitudes of the atoms around their equilibrium positions (B-factors) in the crystal structures, was also used. B-factors encompass a lot of information on the overall biomolecule structure; for example, these parameters depend on conformational disorder, thermal motion paths, and the rotameric state of amino acids side-chains. B-factors also show dependence on the three-dimensional structure as well as protein flexibility (Kanapeckaitė et al., 2021; Faraggi et al., 2009). Normalised dihedral angles (standard deviation scaling to account for variability and distribution) and scaled B-factors (min-max scaling) (Eq.1) were integrated into the Fi-score equation (Eq.2). It is important to highlight that B-factors need to be scaled so that different structural files can be compared and the dihedral angle normalisation transforms angular data into adjusted values based on the overall variability (Kanapeckaitė; et al., 2021).

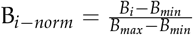

**Equation 1.** Min-max normalisation and scaling of B-factor where *B_i–norm_* is a scaled B-factor, *B_i_* -B-factor for a selected *C_α_* atom in a chain, Bmax - the largest B-factor value for all *C_α_* B-factors in a protein, Bmin - the smallest B-factor value for all *C_α_* B-factors in a protein. B-factor normalisation is based on the full length protein.

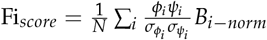

**Equation 2.** Fi-score evaluation where N is the total number of atoms for which dihedral angle information is available, *ϕ* and *ψ* values represent dihedral angles for a specific *C_α_* atom, *σ_ϕi_* and *σ_ψi_* represent corresponding standard deviations for the torsion angles and *B_i–norm_* is a normalised B-factor value for the *C_α_* atom. B-factor, *σ_ϕi_* and *σ_ψi_* normalisation is based on the full length protein.

In order to identify meaningful clusters based on the structural complexity, Gaussian mixture models (GMM) were selected as a primary machine learning classifier (Kanapeckaitė et al., 2021). The strength of GMM lies in the probabilistic model nature since all data points are assumed to be derived from a mixture of a finite number of Gaussian distributions with unknown parameters (Kanapeckaitė et al., 2021; Reynolds, 2015). Consequently, the soft classification of GMM where a data point has a probability of belonging to a cluster is much more suitable to assess biological parameters compared to other hard classification techniques in machine learning, such as k-means, which provide only a strict separation between classes. GMM pipeline offers a number of benefits to categorise protein structural features and the information can be used to explore amino acid grouping based on their physicochemical parameters. The designed GMM implementation takes care of the information criterion assessment to fine tune the number of clusters for modelling and predicts the best suited model for the expectation-maximisation (EM) algorithm to maximise the likelihood of data point assignments (Kanapeckaitė et al., 2021; Reynolds, 2015).

Nur77 protein was used as a case example to demonstrate various package functionalities. Nuclear receptor subfamily 4 group A member 1 (NR4A1), also known as Nur77/TR3/NGFIB, is a member of the nuclear receptor superfamily and regulates the expression of multiple target genes (Wu and Chen, 2018). This nuclear receptor is classified as an orphan receptor since there are no known endogenous ligands. Nur77 has the typical structure of a nuclear receptor which consists of an N-terminal domain, a DNA binding domain, and a ligand-binding domain. This regulatory protein plays many potentially therapeutically relevant roles regulating cell proliferation and apoptosis (Wu and Chen, 2018). Consequently, the Nur77 protein is an excellent example to highlight how in-depth structural analysis and classification could be beneficial in better understanding protein functions and finding potentially druggable binding sites or identifying ligands.

Based on the need to develop integratable and specialised tools for protein analyses, the *Fiscore* package was developed to assist with a wide spectrum of research questions ranging from exploratory analyses to therapeutic target assessment (Fig. 1) The introduced set of new tools provides an interactive exploration of targets with an easy integration into downstream analyses. Importantly, the package and associated tools are written to be easy to use and facilitate analyses for non-specialists in structural biology or machine learning.

**Figure 1:**
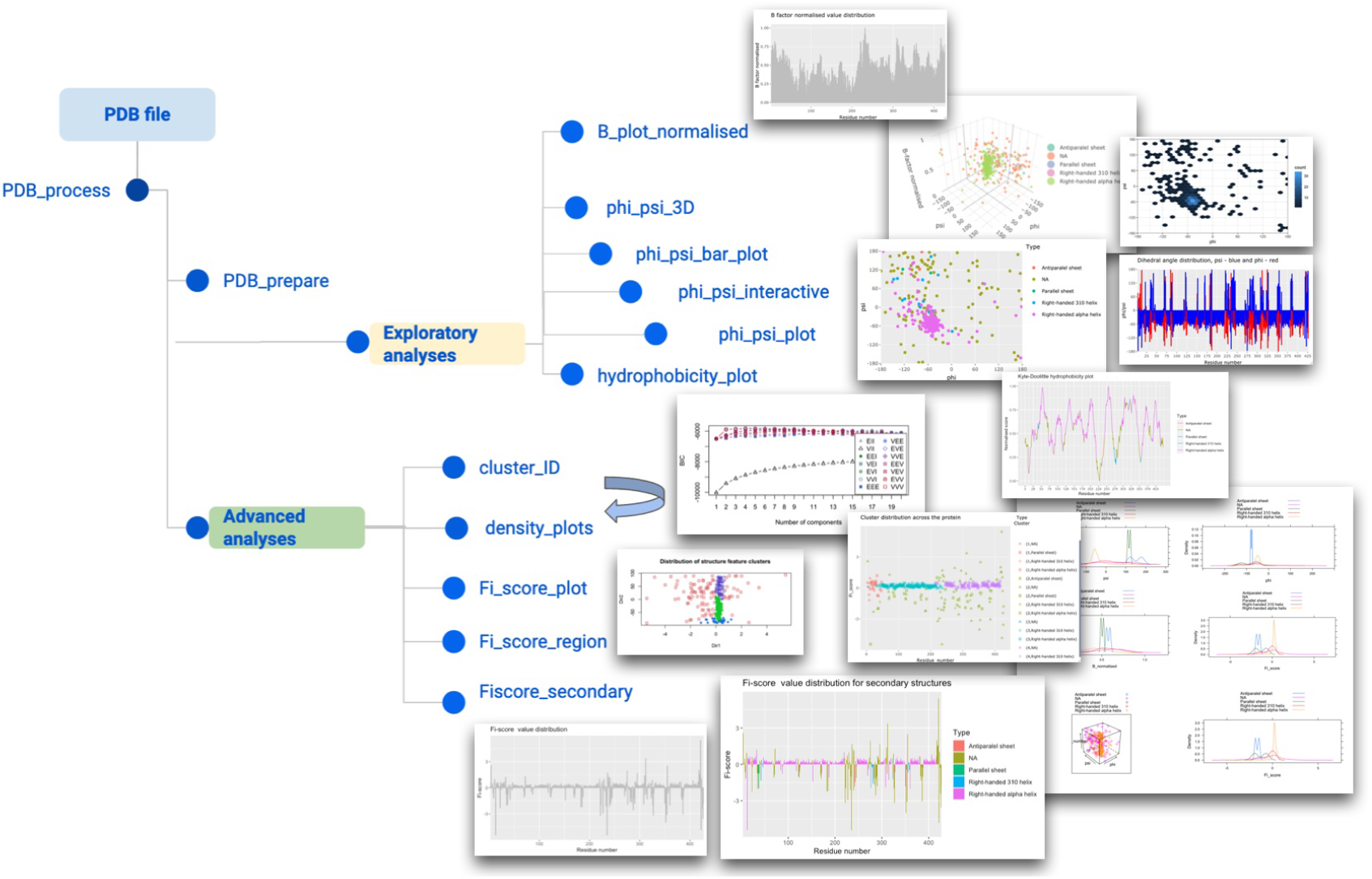
Schematic visualisation of the package features.

## Methods

*Fiscore* package architecture is divided into exploratory and advanced functions (Fig. 1). Several key packages, such as ggplot2 (Wickham, 2016), Bio3D (B.J. et al., 2006), plotly (Sievert, 2020), and mclust (Scrucca et al., 2016), are also employed to create an easy-to-use analytical environment where a user-friendly machine learning pipeline of GMM (Kanapeckaitė et al., 2021) allows for a robust structural analysis. GMM implementation is designed to include the optimal cluster number evaluation (Bayesian information criterion; BIC), automatic model fitting in the EM phase of clustering, model-based hierarchical clustering, density estimation, as well as discriminant analysis (Kanapeckaitė et al., 2021; Kanapeckaite, 2021). Researchers also have an option to perform advanced exploratory studies or integrate the package into their development pipelines. *Fiscore* also takes care of raw data pre-processing and evaluation with optional settings to adjust how the analyses are performed. The package was built using functional programming principles with several R S3 methods to create objects for PDB files (Chambers, 2014). *Fiscore* is accompanied by documentation and vignette files to help the users with their analyses (Kanapeckaite, 2021). Since PDB files are typically large, the documentation provides a compressed testing environment as well as a detailed tutorial. Additional visualisations were generated with PyMol (DeLano, 2021). Proteins were retrieved from Protein Data Bank database (Berman et al.). Protein sequence alignments were performed with PSI-BLAST using default parameters and a single iteration (Altschul et al., 1997). Hydrophobicity plots for Nur77 functional analysis were generated with the following parameters: window = 15,weight = 25, model=“exponential”. Student t-test (two-sided, unpaired, sig. level=95%) was performed in R programming environment.

## Results

### Data preparation

The workflow begins with the PDB file pre-processing and preparation. The user should also generally assess if the structure is suitable for the analysis; that is, the crystallographic data provides a good resolution and there are no or a minimal number of breakages within the reported structure. Function *PDB_process* takes a PDB file name which can be expressed as ‘6KZ5.pdb’ or ‘path/to/the/file/6KZ5.pdb’. One of the function’s dependencies is package Bio3D (B.J. et al., 2006), this useful package provides several tools to begin any PDB file analysis. In addition, the *PDB_process* function can take a ‘path’ parameter which can point to a directory where to split PDB files into separate chain files (necessary for the downstream analysis). If this option is left empty, a folder in the working directory will be created automatically. If the user splits multiple PDB files in a loop, they will be continuously added to the same folder. After the processing, the function *PDB_process* returns a list of split chain names. It is important to highlight that for the downstream processing PDB files need to be split so that separate chains can be analysed independently.

After a file or files are pre-processed the function *PDB_prepare* can be used to prepare a PDB file to generate Fi-score and normalised B-factor values as well as secondary structure designations. The function takes a PDB file name that was split into separate chains, e.g. ‘6KZ5_A.pdb’, where a letter designates a split chain. The file is then cleaned and only the complete entries for amino acids are kept for the analysis, i.e. amino acids from the terminal residues that do not contain both dihedral angles are removed. The function returns a data frame with protein secondary structure information ‘*Type*’, Fi-score values per residue ‘*Fi_score*’, as well as normalised B-factor values for each amino acid *C_α_* ‘*B_normalised*’ (Fig. 2). Extracting protein secondary structure information, i.e. ‘*Type*’, helps to prepare a data object so that the information about a target could be supplied into cheminformatics or other bioinformatics pipelines where structural features are important to assess protein sites and amino acid composition.

**Figure 2:**
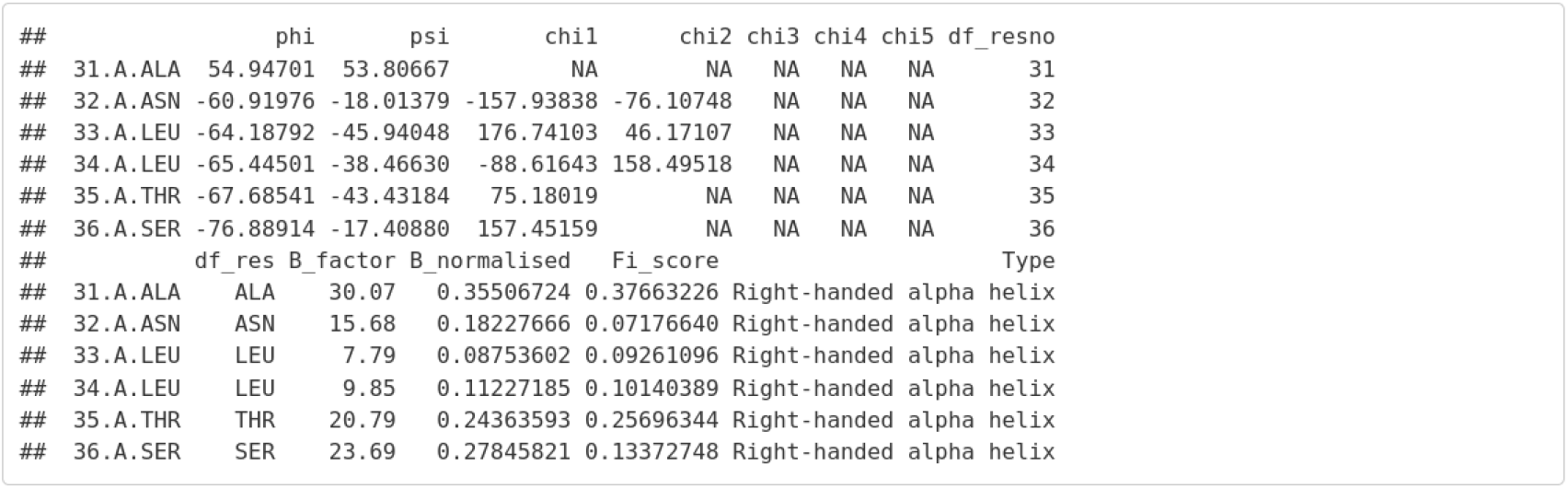
PDB file processing output.

Function calls are simple and user-friendly:

~~~
#General function for pre-processing raw PDB files
pdb_df<-PDB_process(pdb_path)
#Cleaning and preparation of PDB file
pdb_df<-PDB_prepare(pdb_path)
#Explore the output
head(pdb_df)
#The package allows to call test data directly for the Nur77 example file
pdb_path<- system.file(“extdata”, “6kz5.pdb”, package=“Fiscore”)
~~~

### Exploratory analyses

The scope of the exploratory analyses provides options to evaluate physicochemical parameters, such as dihedral angles, B-factors, or hydrophobicity scores, and visualise their distribution (Fig. 1).

Basic analyses are accessed through simple function calls to explore how dihedral angles and B-factors are distributed. These analyses offer interactive and easy visualisations of key parameters that are currently not offered in any other package.

~~~
#Calling a Ramachandran plot function
phi_psi_plot(pdb_df)
#Visualisation of dihedral angle juxtaposed distributions
phi_psi_bar_plot(pdb_df)
#B plot value visualisation
B_plot_normalised(pdb_df)
#Interactive plot to map amino acids via 2D distribution
#to precisely see what parameters an individual amino acid has
phi_psi_interactive(pdb_df)
#3D visualisation of dihedral angles and B-factor values
phi_psi_3D(pdb_df)
~~~

Especially useful functionality is the hydrophobicity visualisation with the superimposed secondary structure elements. To the author’s knowledge, there are currently no tools implementing such a visualisation (Fig. 3). The package provides an easy to use wrapper:

~~~
hydrophobicity_plot(pdb_df,window = 9,weight = 25,model = “linear”)
#Alternatively an exponential model can be selected
hydrophobicity_plot(pdb_df,window = 9,weight = 25,model = “exponential”)
~~~

**Figure 3:**
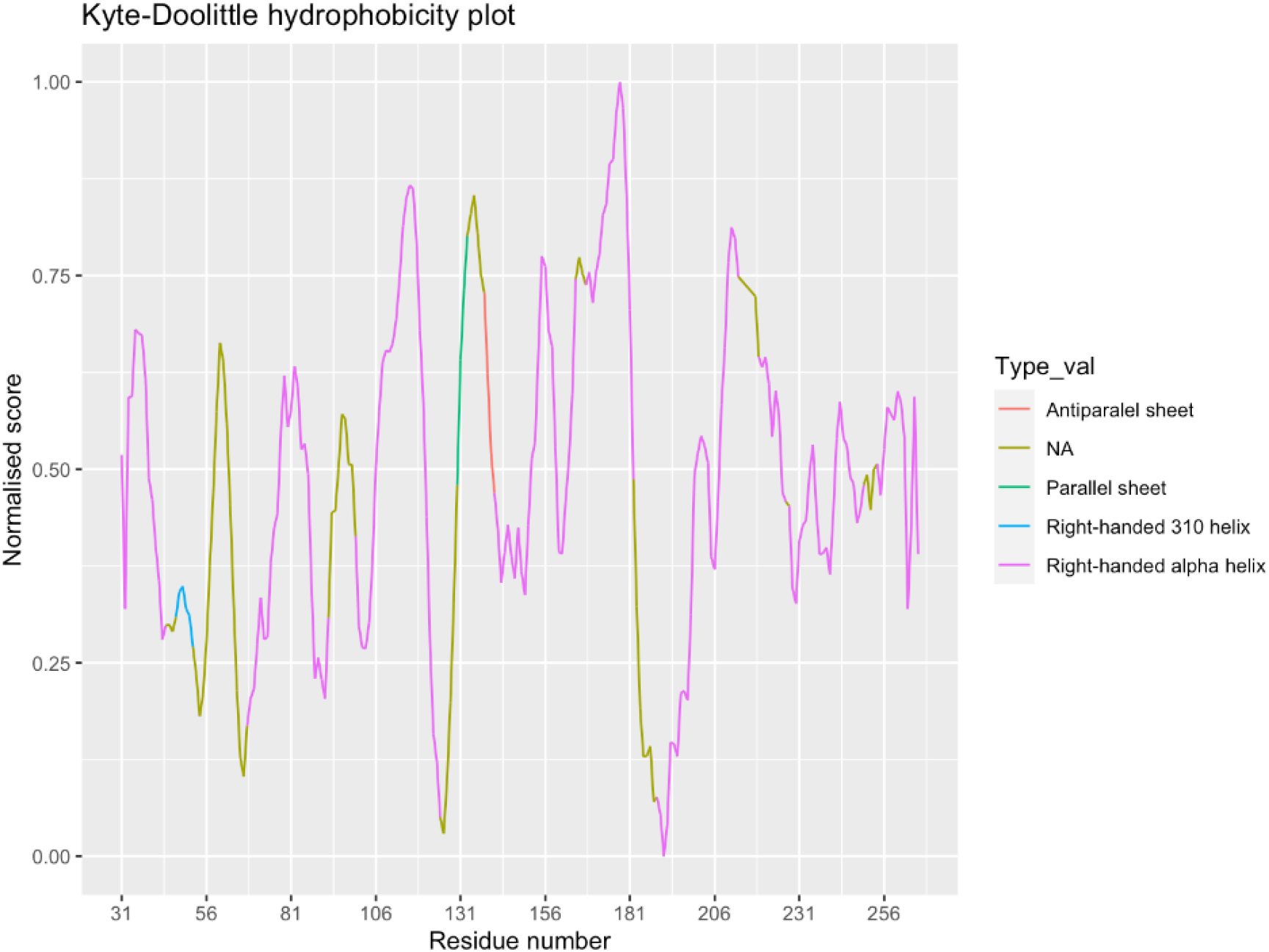
Hydrophobicity plot with secondary structure superimposition.

Employing the introduced hydrophobicity analysis to assess the nuclear receptor reveals an overall dynamic profile for the protein. Moreover, Nur77 evidently contains a relatively large number of right-handed alpha helices with the majority showing a hydrophobic profile, i.e. the larger the score, the more hydrophobic the region. Some likely disordered regions can be seen spanning 50-70 amino acids (Fig. 4). Another interesting region is around 126-136 amino acids since these amino acids undergo significant shifts in their hydrophilicity and hydrophobicity. Similarly, the region around 180-210 amino acids appears to be actively changing preferences from little solvent to being solvent exposed. This might suggest that the site undergoes considerable movements or actively engages other proteins or the DNA sequence. The disordered elements in this sequence stretch also imply that the region has to likely accommodate various rearrangements. Thus, studying these sites could provide hints at functionally important protein domains or subdomains (Fig. 3&4). Finally, evaluating N and C terminal sites for the purpose of protein engineering, we can see that a histidine tag would not significantly disrupt the conformation of the molecule and the C-terminus is probably the best site for the tag.

**Figure 4:**
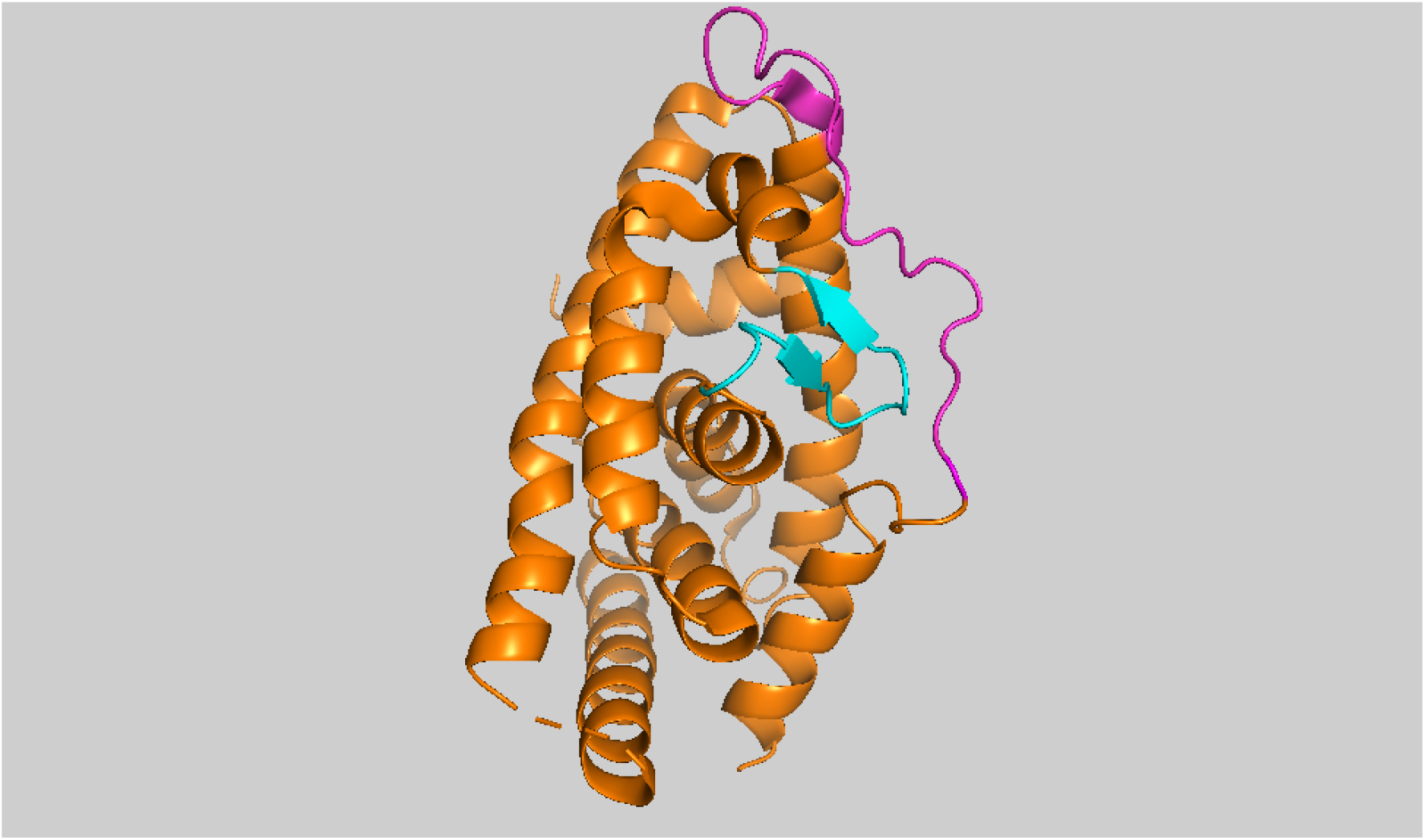
Nur77 protein where magenta highlights are used to define a likely disordered region between 50 and 70 amino acids and cyan indicates a region between 127 and 140 amino acids.

It is worth commenting on the derivation of the hydrophobicity scoring since the algorithmic nature of the process provides several important analytical angles. The function builds on the Kyte-Doolittle hydrophobicity scale (Kanapeckaitė et al., 2021; Kyte and Doolittle, 1982) to detect hydrophobic regions in proteins. Regions with a positive value are hydrophobic and those with negative values are hydrophilic. This scale can be used to identify both surface-exposed as well as transmembrane regions, depending on the window size used. However, to make comparisons easier, the original scale is transformed from 0 to 1. The function requires a PDB data frame generated by *PDB_prepare* and the user needs to specify a ‘window’ parameter to determine the size of a window for hydrophobicity calculations. The selection must be any odd number between 3 and 21 with the default being 21. Another parameter is ‘weight’ that needs to be supplied to the function to establish a relative weight of the window edges compared to the window center (%); the default setting is 100%. Finally, a ‘model’ parameter provides an option for weight calculation; that is, the selection determines whether weights are calculated linearly (*y* = *k* · *x* + *b*) or exponentially (*y* = *a* · *b^x^*); the default model is ‘linear’. The function evaluates each amino acid in a selected window where a hydrophobic influence from the surrounding amino acids is calculated in. While the terminal amino acids cannot be included into the window for centering and weighing, they are assigned unweighted values based on the Kyte-Doolittle scale (Kyte and Doolittle, 1982). The plot values are all scaled from 0 to 1 so that different proteins can be compared without the need to convert.

Thus, the hydrophobicity analysis can be especially useful when preparing to engineer proteins for various expression systems as the superimposition of structural features and hydrophobicity scores can help deciding if a protein region or domain is likely to be solvent exposed or prefer hydrophobic environments. For example, assessing the N or C terminal amino acid hydrophobicity and structural milieu can help selecting which terminal should be tagged (as was demonstrated with Nur77). Moreover, this tool could be broadly applied in drug discovery studies involving the assessment of protein-protein interactions, protein-nucleic acid interactions, and membrane association events.

### Advanced analyses

Advanced analyses provide an opportunity to evaluate Fi-score distributions and take advantage of a streamlined GMM pipeline. The main impetus for the development of this pipeline was the need for functions and data modelling tools that could be made accessible to non-experts. More advanced users can supply custom parameters for the GMM workflow and extract probabilities from the output to use scores in other analyses or integrate the values in their own discovery pipelines.

~~~
#Fi-score distribution plot to explore scores for corresponding amino acids
Fi_score_plot(pdb_df)
#Fi-score for a selected region
#this value for multiple sites can be stored in relational databases
Fi_score_region(pdb_df,50,70)
#Plot of Fi-score values with superimposed secondary structures
Fiscore_secondary(pdb_df)
~~~

For example, a Fi-score distribution plot captures several interesting regions in Nur77 around the 50, 130, and 180 amino acids (Fig. 5). Some other regions are also picked up which should be studied in more detail based on the amino acid composition and 3D conformations. The uncovered characteristics can be juxtaposed to other similar sites to better understand interaction mechanisms. Such approaches are especially useful when comparing known structures with newly identified ones.

**Figure 5:**
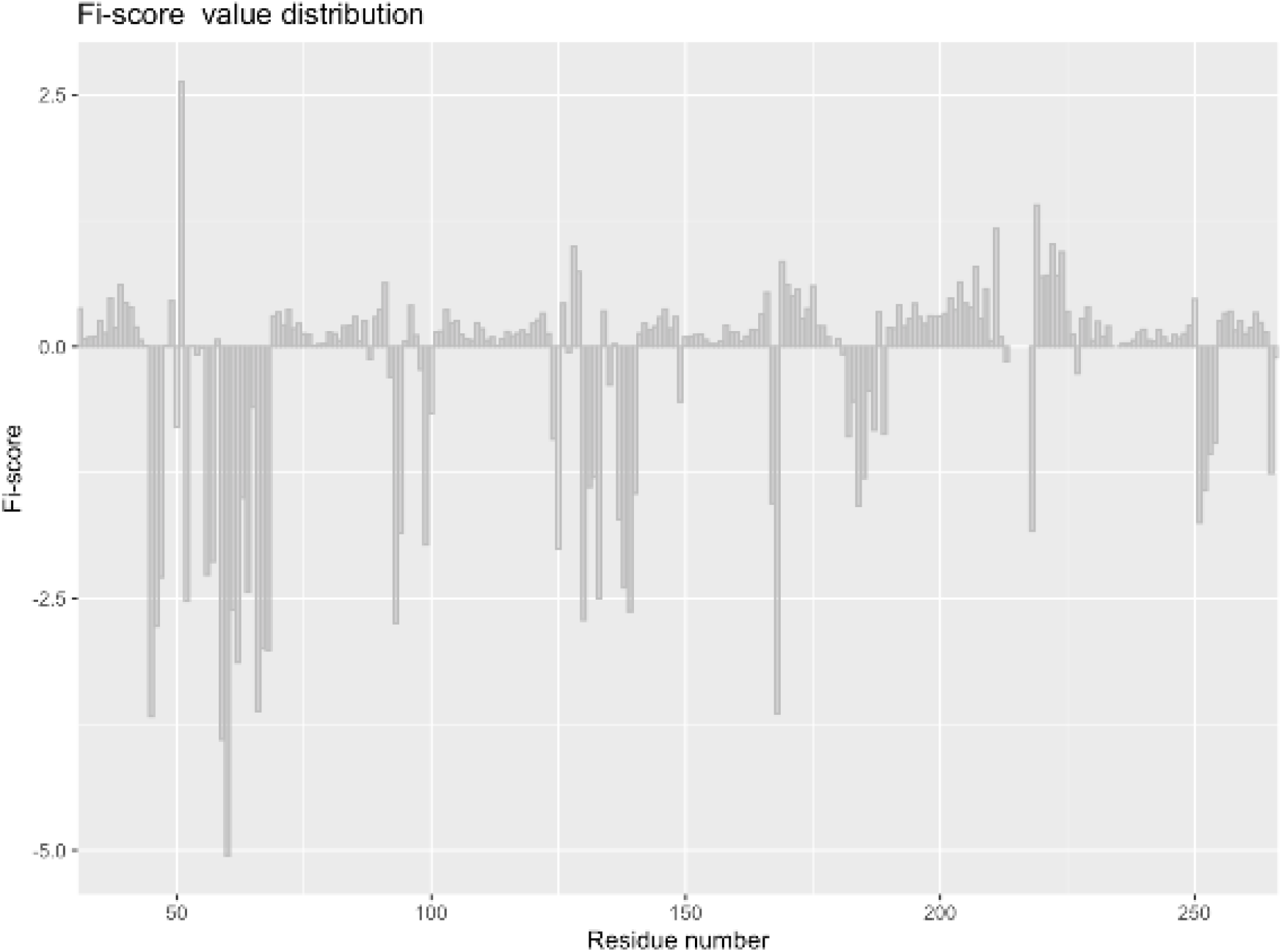
Fi-score distribution for Nur77.

Extracted Fi-score values can be used in machine learning modelling and this is enabled through the function *cluster_ID*. This function groups structural features using the Fi-score and Gaussian mixture modelling where an optimal number of clusters and a model to be fitted during the EM phase of clustering for GMM are automatically selected (Fig. 6). The output of this analytical tool summarises cluster information and also provides plots to visualise the identified clusters based on the cluster number and BIC value (Fig. 7).

~~~
df<-cluster_ID(pdb_df)
#User selected parameters
df<-cluster_ID(pdb_df,clusters = 5, modelNames = “VVI”)
~~~

**Figure 6:**
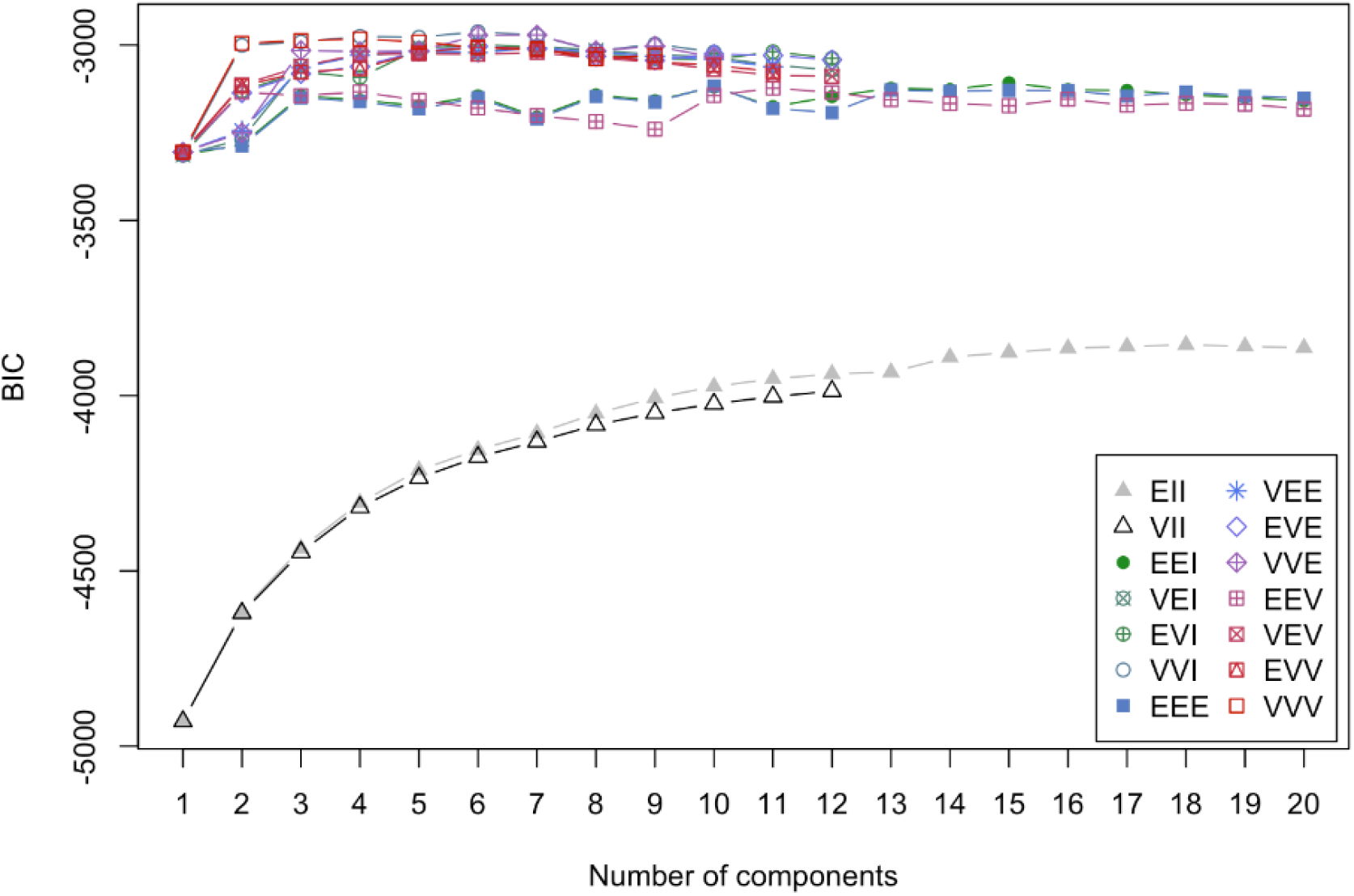
Gaussian mixture modelling output showing Bayesian information criterion evaluation.

**Figure 7:**
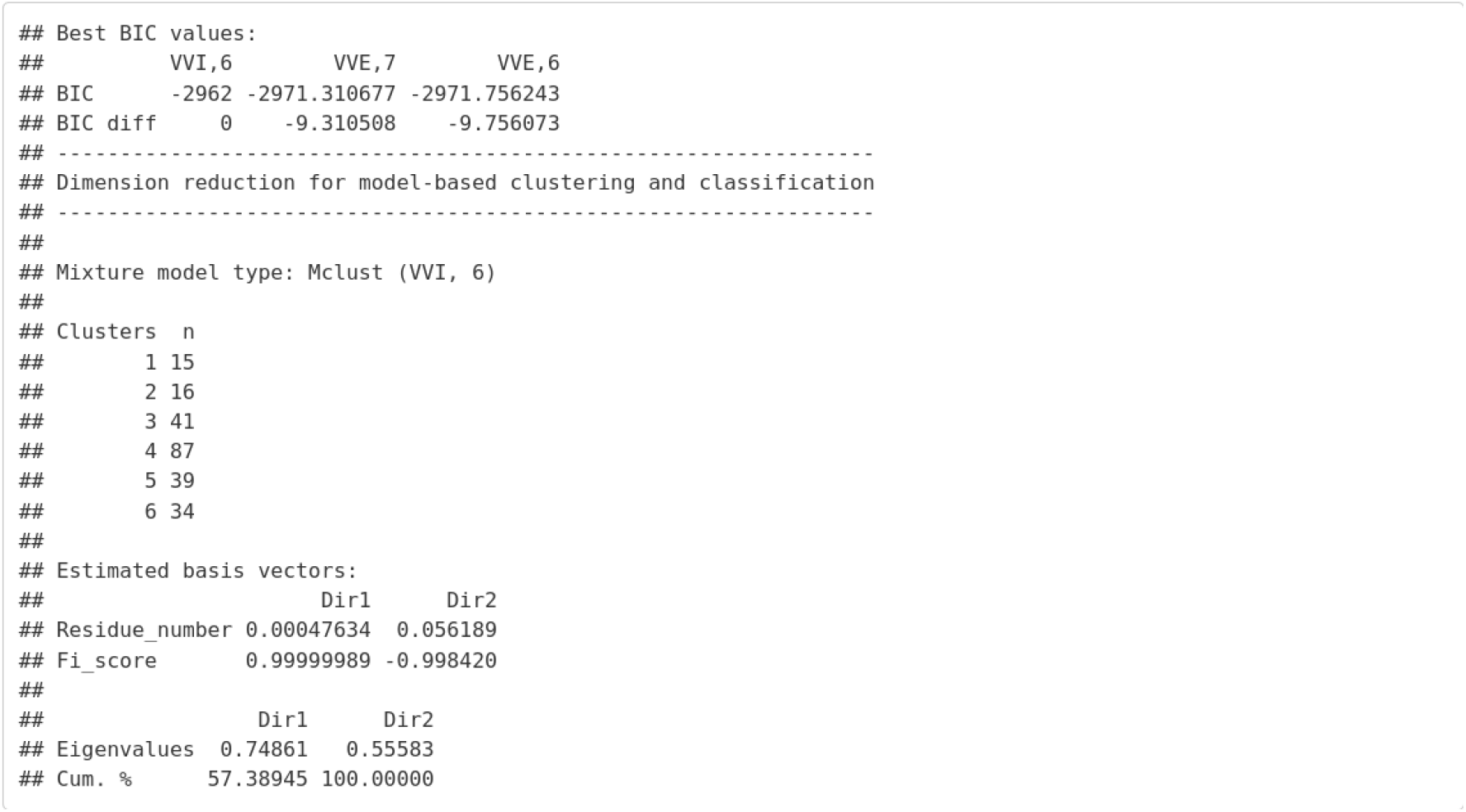
Output table for Gaussian mixture modelling evaluation.

The users are advised to set seed for more reproducible results when initiating their projects. *cluster_ID* takes a data frame containing a processed PDB file with Fi-score values as well as a number of clusters to consider during model selection; by default 20 clusters (‘max_range’) are explored. In addition, a ‘secondary_structures’ parameter is needed to define whether the information on secondary structure elements from the PDB file needs to be included when plotting; the default value is TRUE. Researchers also have an option to select a cluster number to test ‘clusters’ together with ‘modelNames’. However, it is important to stress that both optional entries need to be selected and defined, if the user wants to test other clustering options that were not provided by the automated BIC output. This is an advanced option and the user should assess the BIC output to decide which model and what cluster number he or she wants to try out.

A dimension reduction method for the visualisation of clustering is also automatically provided (Fig. 10). Dimension reduction is a useful technique to explore multi-dimensional biological data through key eigenvalues that define the largest information content of the explored features (Scrucca et al., 2016). In other words, one can infer how well the explored characteristics define the data and if the classification is sufficient for downstream analyses. For example, in the case of Nur77 Fi-score clustering, this analysis allowed assessing how well the number of clusters separates data points based on their distribution features. Nur77 has six clusters which might indicate functionally and structurally distinct regions in the target protein. It appears that the data points are well separated into groups accounting for the different variability. The dimension reduction approach could also help deciding if a different number of clusters might better classify Fi-scores.

In addition, one of the most valuable features of this set of functions is to generate clusters with secondary structure information (Fig. 8&9).The produced interactive plots enable researchers to explore structural characteristics of a protein of interest (Fig. 8&9). Thus, the subdivision of a protein structure based on the physicochemical features offers a new way to detect and explore functional sites or structural elements. Figures 9 and 10 clearly indicate that some structural elements in Nur77 are likely similar in their function and physicochemical characteristics. For example, different types of helices as well as beta sheets in some cases overlap in their Fi-score characteristics and the assigned cluster type. This detailed capture of structural elements can help evaluate conformational outliers or infer similarities for different motifs. Moreover, it can be clearly seen that the region around the 50 amino acid is set to be distinct from the other two sites around 130 and 180 amino acids which could suggest overall different motion and interaction profiles. These findings also correspond to the earlier observations for the hydrophobicity features (Fig. 3). A similar trend can be seen for N and C terminus clusters which form distinct groups and might indicate sites where the receptor mediates specific functions (Wu and Chen, 2018).

**Figure 8:**
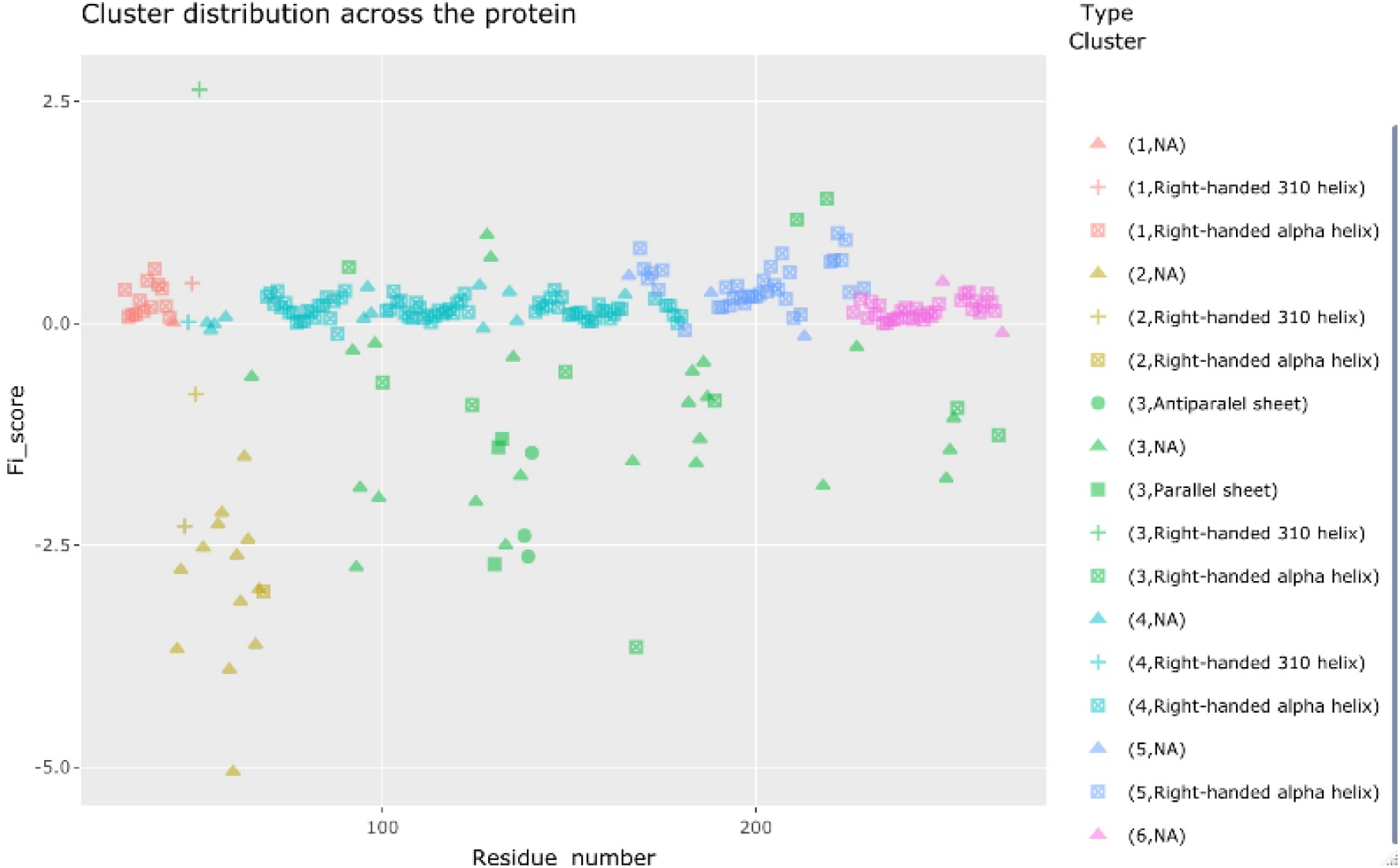
The Nur77 protein cluster identification with secondary structure elements.

**Figure 9:**
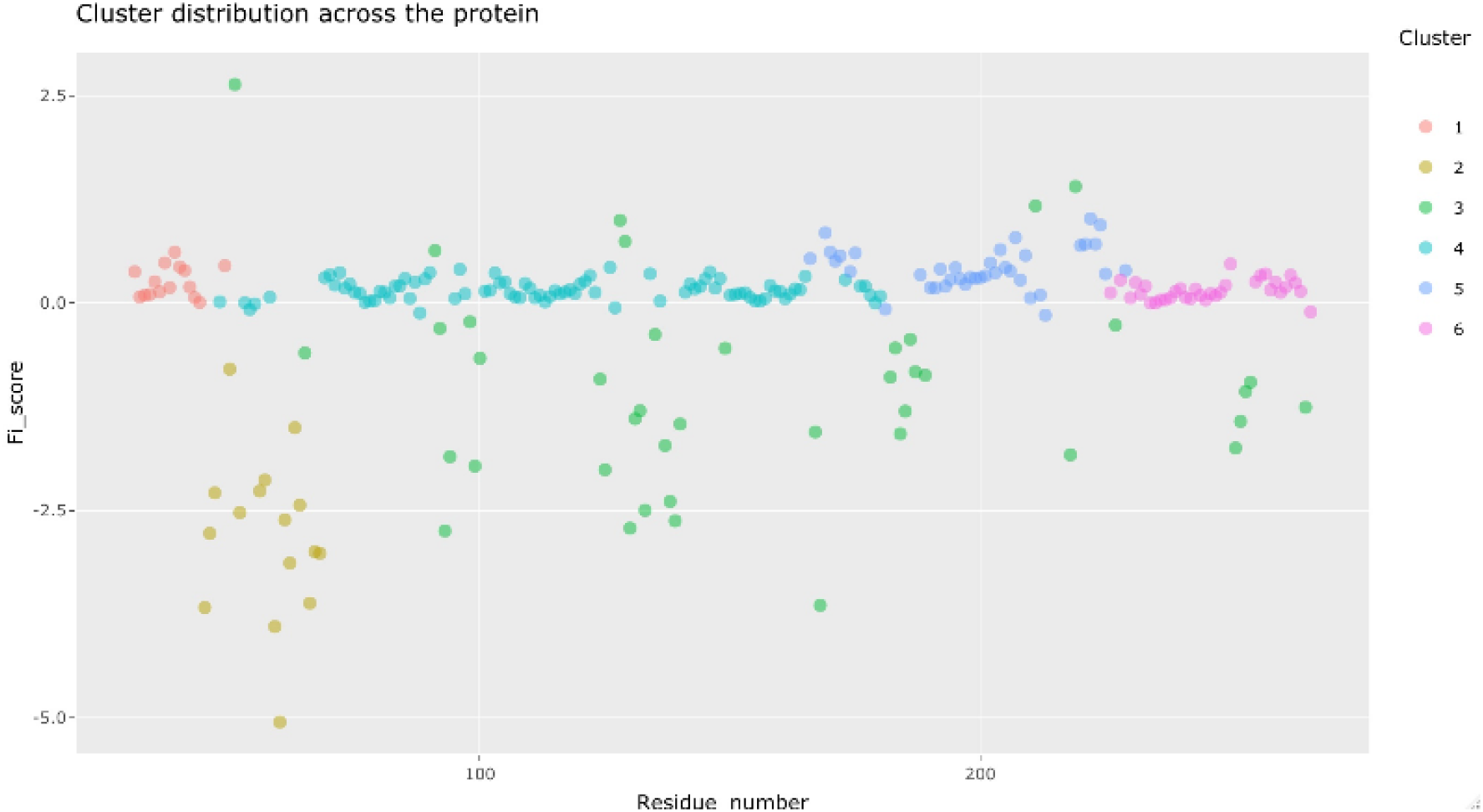
Nur77 cluster identification.

**Figure 10:**
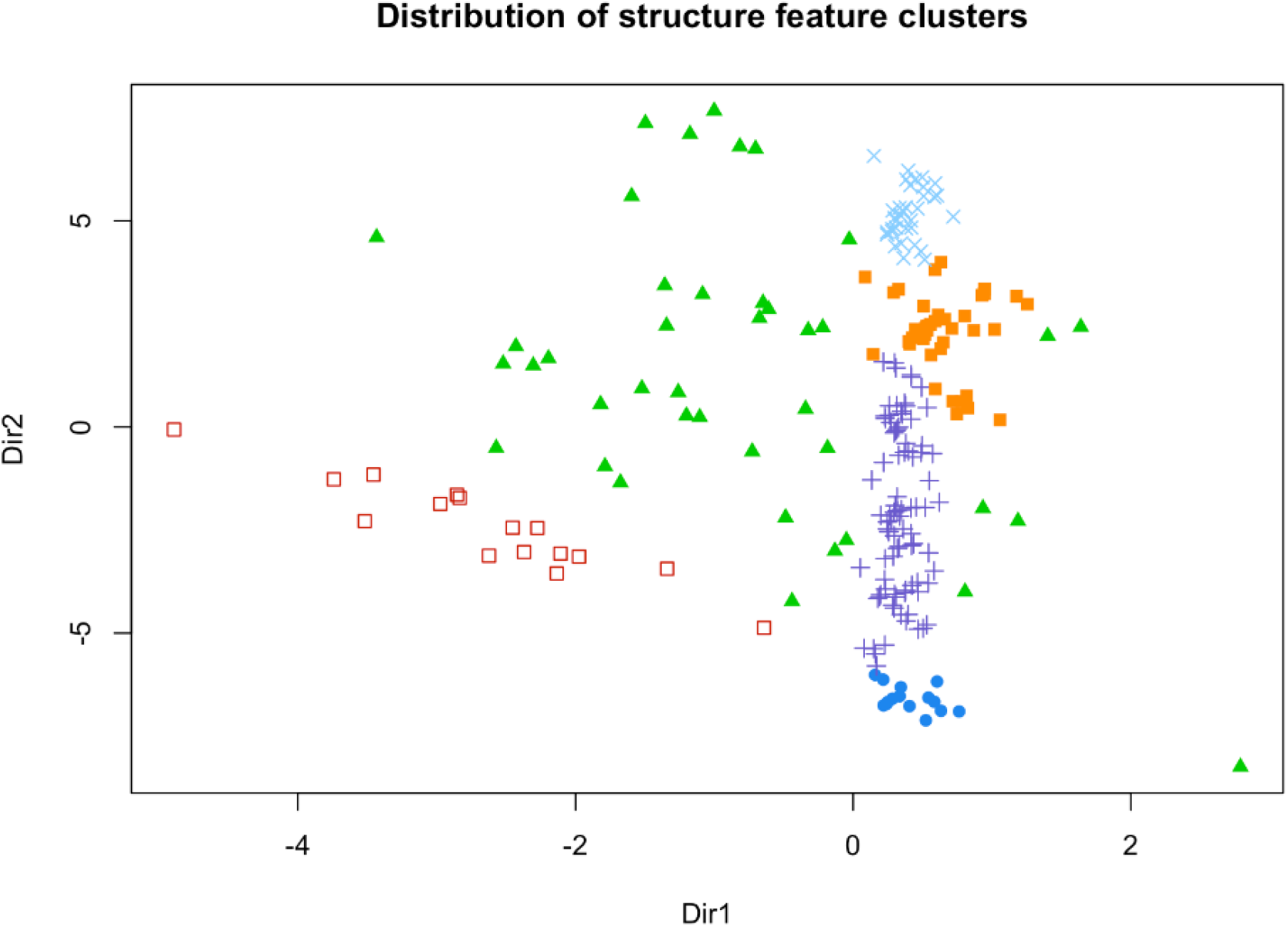
Dimension reduction plot for the identified clusters.

All previous analyses tie in with the function *density_plots* which provides a density plot set for *ϕ*/*ψ* angle distributions, Fi-score, and normalised B-factor. 3D visualisation of dihedral angle distribution for every residue is also included. The plots can be used for a quick assessment of the overall parameters as well as to summarise the observations. Density plots are optimal to evaluate how well the selected features or scores separate protein structural elements and if, for example, a protein structure is of good quality (i.e., dihedral angles, B-factors, or Fi-scores provide reasonable separation between elements). The function also gives another reference point to establish if the selected number of clusters differentiates residues well based on the secondary structure elements. In order to get this information, the user is only required to supply the output from the *cluster_ID* function (Fig. 11).

**Figure 11:**
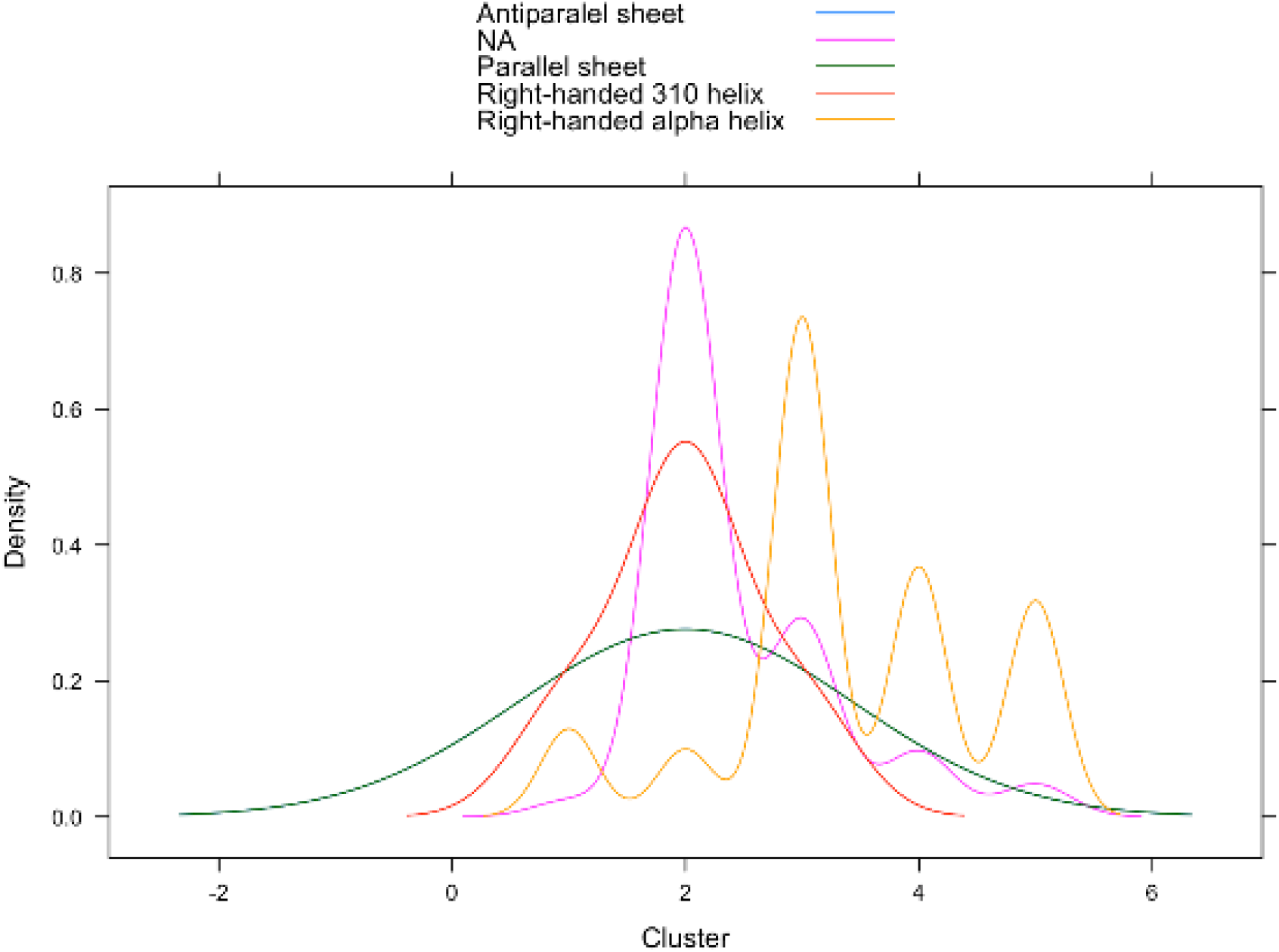
Protein cluster density plots.

~~~
# Data summary and evaluation
density_plots(pdb_df)
# Data summary and evaluation including GMM outputs
cluster_IDs<-cluster_ID(pdb_df)
density_plots(cluster_IDs)
~~~

### Case study: exploring potential ligands for the Nur77 orphan receptor

To demonstrate some *Fiscore* applications, potential ligands were searched for the Nur77 receptor. The first analysis step involved searching for other similar human proteins that did not belong to the nuclear receptor subfamily 4. PSI-BLAST alignment analysis led to several candidate proteins, namely retinoic acid receptor alpha (PDB ID: 1FBY) and estrogen-related receptor gamma (PDB ID: 6KNR) (Altschul et al., 1997). These proteins showed a good alignment to the Nur77 ligand binding domain sequence (average percent identity 31.68 %; Suppl. Table 1) and were subsequently used for structural and functional exploration. Comparing Nur77 Fi-scores with the retinoic acid receptor alpha and estrogen-related receptor gamma Fi-score distributions revealed several interesting patterns (Fig. 12). The shaded blue region highlights a matching distribution pattern for all the proteins and Student t-test confirmed that none of the distributions differed significantly (Fig. 12; Suppl. Table 2). Intriguingly, this region is involved in mediating interaction with retinoic acid in the retinoic acid receptor alpha (Fig. 13). Moreover, despite different amino acid composition, the key physicochemical features are preserved in this site across the investigated proteins as can be seen from the superimposition studies (Fig. 13). This suggests an interesting possibility that Nur77 with no known ligands might bind to chemical entities similar to retinoic acid (Wu and Chen, 2018). This is also supported by the alignment data and hydrophobicity plots (Suppl Fig. 1-3) where Nur77 and the retinoic acid receptor alpha show substantial structural and physicochemical overlaps for this interactor site. Thus, these examples reveal that extracting patterns for database search could help with identifying proteins that have shared features without necessarily performing multiple alignments and visual inspections of the structures. These analytical principles can also be applied to explore other proteins of interest and their potential ligands.

**Figure 12:**
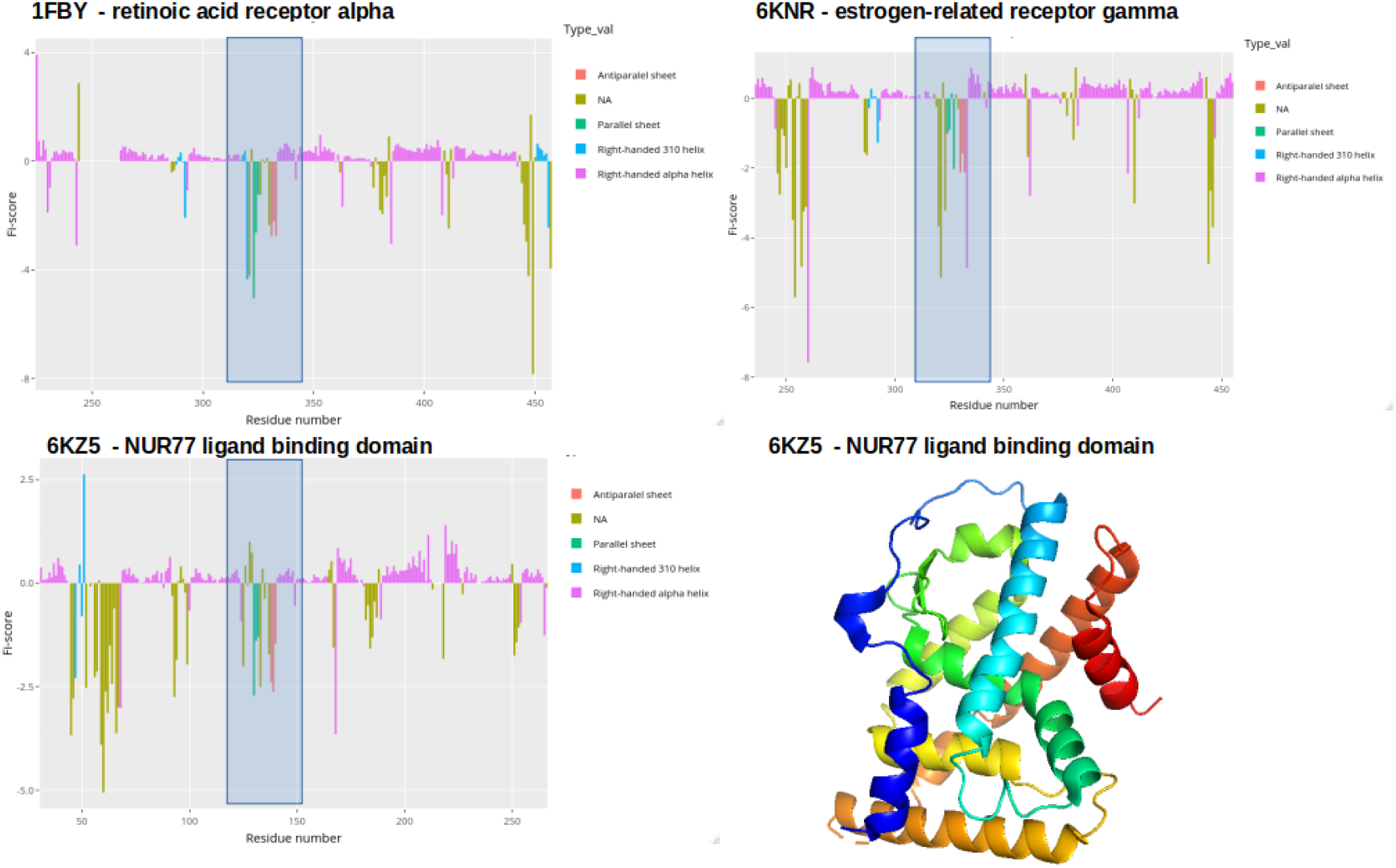
Fi-score distribution plots with Nur77 ligand binding domain (PDB ID:6KZ5), Retinoic acid receptor alpha (PDB ID: 1FBY), and estrogen-related receptor gamma (PDB ID: 6KNR). Rainbow spectrum of the Nur77 structure allows to visualise the sequence from N-terminal (blue) to C-terminal (red).

**Figure 13:**
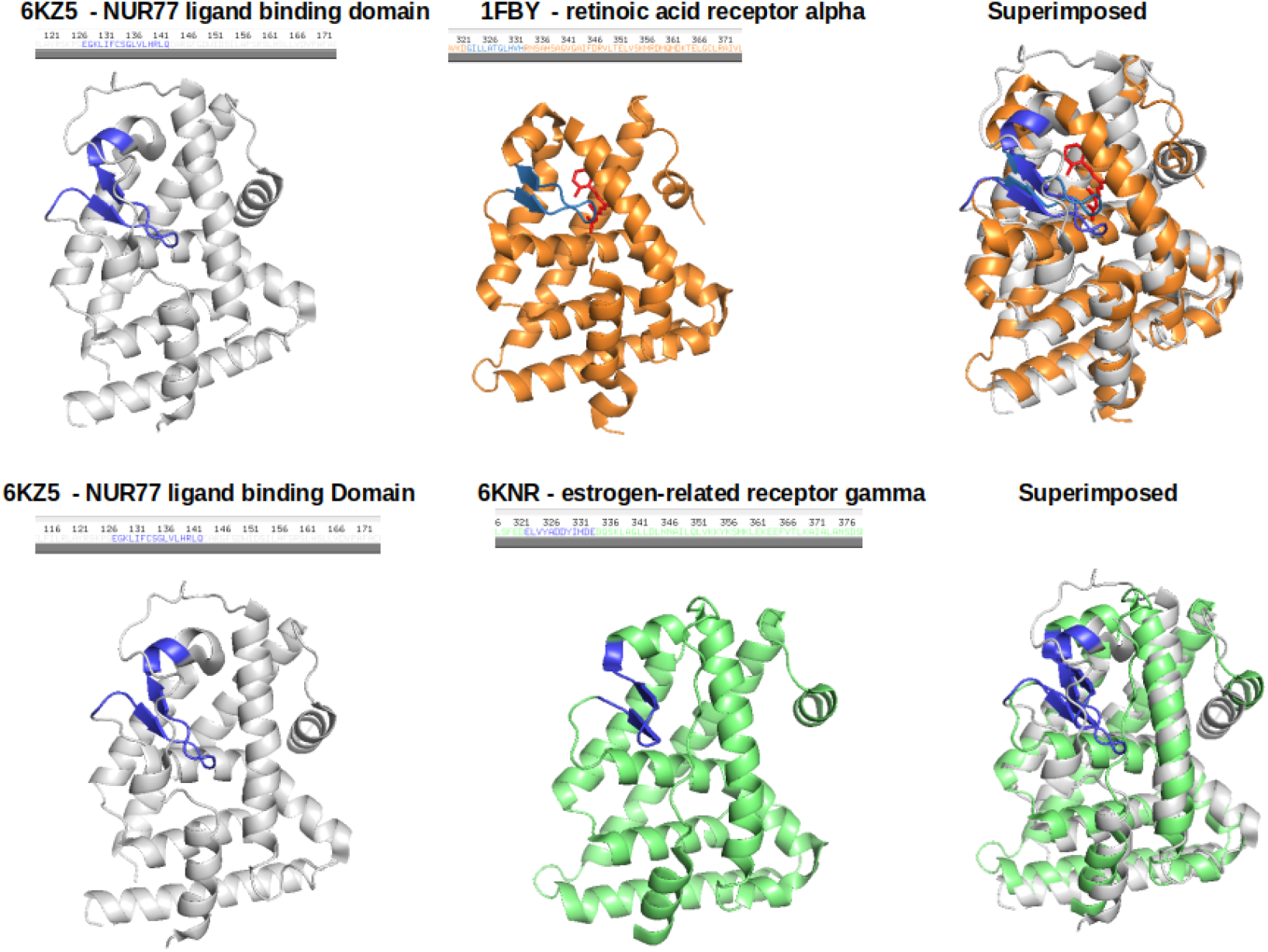
PyMol generated plots to visualise protein structures where blue colors indicate the region identified through Fi-score patterns where cis-9 retinoic acid is highlighted in red.

## Discussion

*Fiscore* package (Fig. 1) allows a user-friendly exploration of PDB structural data and integration with various machine learning methods. The package was benchmarked through several analytical stages that involved a diverse set of proteins (3352) to assess scoring principles (Kanapeckaitė; et al., 2021) and package functionalities (1337 structures) (Kanapeckaite, 2021). With a number of helpful functions, including distribution analyses or hydrophobicity assessment in the context of structural elements, *Fiscore* enables the exploration of new target families and comprehensive data integration since the described fingerprinting captures protein sequence and physicochemical properties. Such analyses could be very helpful when exploring therapeutically relevant proteins. Similarly, *Fiscore* could aid in drug repurposing studies when a chemical compound needs to be juxtaposed to a number of potential targets. This was also demonstrated during a native ligand search for Nur77. In addition, provided tutorials and documentation should guide researchers through their analysis and allow adapting the package based on individual research needs (Kanapeckaitė; et al., 2021). A case study of a selected complex target, the Nur77 nuclear receptor, helped to demonstrate the usefulness of capturing physicochemical data in visual representations. Furthermore, a novel scoring system as well as machine learning applications can lead to interesting insights about sites of structural and functional importance. The retrieved information could be used in comparative studies to search for other proteins that share similar features. Intriguingly, some of the shifts in Fi-score values coincide or precede post-translational modifications in Nur77 (Fig. 5) (Hornbeck et al., 2015) which could be used to classify and fingerprint this data in larger scale exploratory studies.

Another important aspect of the *Fiscore* package is the simplification of complex analytical steps so that the researchers without an extensive background in structural bioinformatics or machine learning could still use the tools for their analyses, such as protein engineering, protein assessment, and data storage based on specific target sites. Thus, the interactive analytical and visualisation tools could become especially relevant in the pharmaceutical industry and drug discovery studies as more complex targets and protein-protein interactions need to be assessed in a streamlined fashion. In other words, ability to translate structural data into parameters could accelerate target classification, target-ligand studies, or machine learning integration. Since target evaluation is paramount for rational therapeutics development, there is an undeniable need for specialised analytical tools and techniques that can be used in R&D or academic research and implementing these novel approaches could significantly improve our ability to assess new targets and develop better therapeutics. As a result, *Fiscore* package was developed to aid with therapeutic target assessment and make machine learning techniques more accessible to a wider scientific audience.

## Acknowledgements

The author would like to thank the anonymous reviewers and code testers for helping to improve the package with their valuable suggestions and advice.

